# On the many advantages of using the VariantExperiment class to store, exchange and analyze SARS-CoV-2 genomic data and associated metadata

**DOI:** 10.1101/2021.04.05.438328

**Authors:** Jérôme Ambroise, Laurent Gatto, Julie Hurel, Bertrand Bearzatto, Jean-Luc Gala

## Abstract

On Friday, 19 March 2021, WHO organized a virtual global workshop highlighting the need for a globally coordinated plan to increase SARS-CoV-2 genetic sequencing capacities to detect SARS-CoV-2 mutations and variants, and to monitor virus genomic evolution worldwide. One week later, in another virtual meeting, it focused on sero epidemiology for SARS-CoV-2 variants of concern and variants of interest. Efficient monitoring of the virus relies on the storage, handling and sharing of the genomic data and the associated metadata. In this manuscript, we demonstrate how the Bioconductor VariantExperiment class addresses these needs, offering a robust and efficient solution to the requirements laid out by the WHO.

## Background

The first COVID-19 case was reported in Wuhan city, Hubei province of China in December 2019. The pathogen causing the disease was soon identified as a novel coronavirus, closely related to severe acute respiratory syndrome coronavirus (SARS-CoV), and renamed novel coronavirus SARS-CoV-2. In light of the growing number of cases at Chinese and around the world, the World Health Organization (WHO) Emergency Committee declared a global health emergency on 30 January 2020.

On Friday, 19 March 2021, WHO organized a virtual global workshop on enhancing sequencing for SARS-CoV-2 in the context of the elaboration of a globally coordinated plan to increase SARS-CoV-2 genetic sequencing capacities to detect SARS-CoV-2 mutations and variants, and to monitor virus genomic evolution worldwide. WHO is indeed working with Member States and partners to increase SARS-CoV-2 sequencing capacities and encourage timely sharing of geographically representative sequences and supporting data. One of the main objectives of the coordinated plan is to promote an efficient discovery and reporting of new Variants of Concern (VOC) among the many existing and upcoming genetic variants within the SARS-CoV-2 genome. During this workshop, the importance of metadata and quality metrics of SARS-CoV-2 genomic data was emphasized. In particular, the lack of current minimum standards of metadata and the need for a system enabling to keep all components well synchronized was emphasized. Being able to store and analyze SARS-COV2 genomic data and associated high-quality metadata will favor the efficient discovery of VOC, Variant of Interest (VOI), or Variant of High Consequences in the upcoming months [1]. Of course, this will be enhanced by sharing of all genomics and associated metadata between countries and continents.

Bioconductor is an open-source, open-development software project for the analysis and comprehension of high-throughput data in genomics and molecular biology. The Bioconductor project aims to enable inter-disciplinary research, collaboration and rapid development of scientific software [2]. Among the packages developed and maintained by the core developers of the project, VariantAnnotation and VariantExperiment can be used to import data from VCF and GDS files and store them into structured data model objects [3,4].

The aim of this paper is to demonstrate the many advantages of using these Bioconductor packages for storing, sharing, and analyzing SARS-CoV-2 genomic data and associated metadata. In order to illustrate this purpose, an object of the VariantExperiment class was created with publicly available SARS-COV-2 genomic (Whole Genome Sequencing WGS) data and used in a R markdown as a demonstration tool.

## Methods

In order to illustrate the advantages of using the VariantExperiment data model, we demonstrate how it can be used to create and manipulate genomics SARS-CoV2 data. To this end, 10 fastq raw data files generated by the MinION Oxford Nanopore Technology (ONT) were downloaded from the European Nucleotide Archive (ENA) repository. Data were mapped to the reference genome MN908947.3 with minimap2 2.17 [5] and SAM files were processed using samtools 1.9. [6] The calling of the mutations was performed using freebayes 1.3.2 [7]. After quality filtering with vcffilter, the resulting VCF file and a dummy metadata (including dummy patients characteristics as well as preanalytical, analytical and bioinformatics parameters) file were processed with the VariantExperiment and the VariantAnnotation packages in order to create the desired object of the VariantExperiment class. Finally, an R markdown report and its html rendering^1^ were created to illustrate the structure and benefit of the object [8].

## Results

The Rmarkdown which illustrates the structure and the manipulation of the VariantExperiment object created from SARS-CoV-2 genomic data is available on github^2^. The structure of the VariantExperiment class is illustrated in Figure 1. The advantages of such Bioconductor objects in the context of SARS-COV2 genomic data analysis are described below.

**Figure 1:**
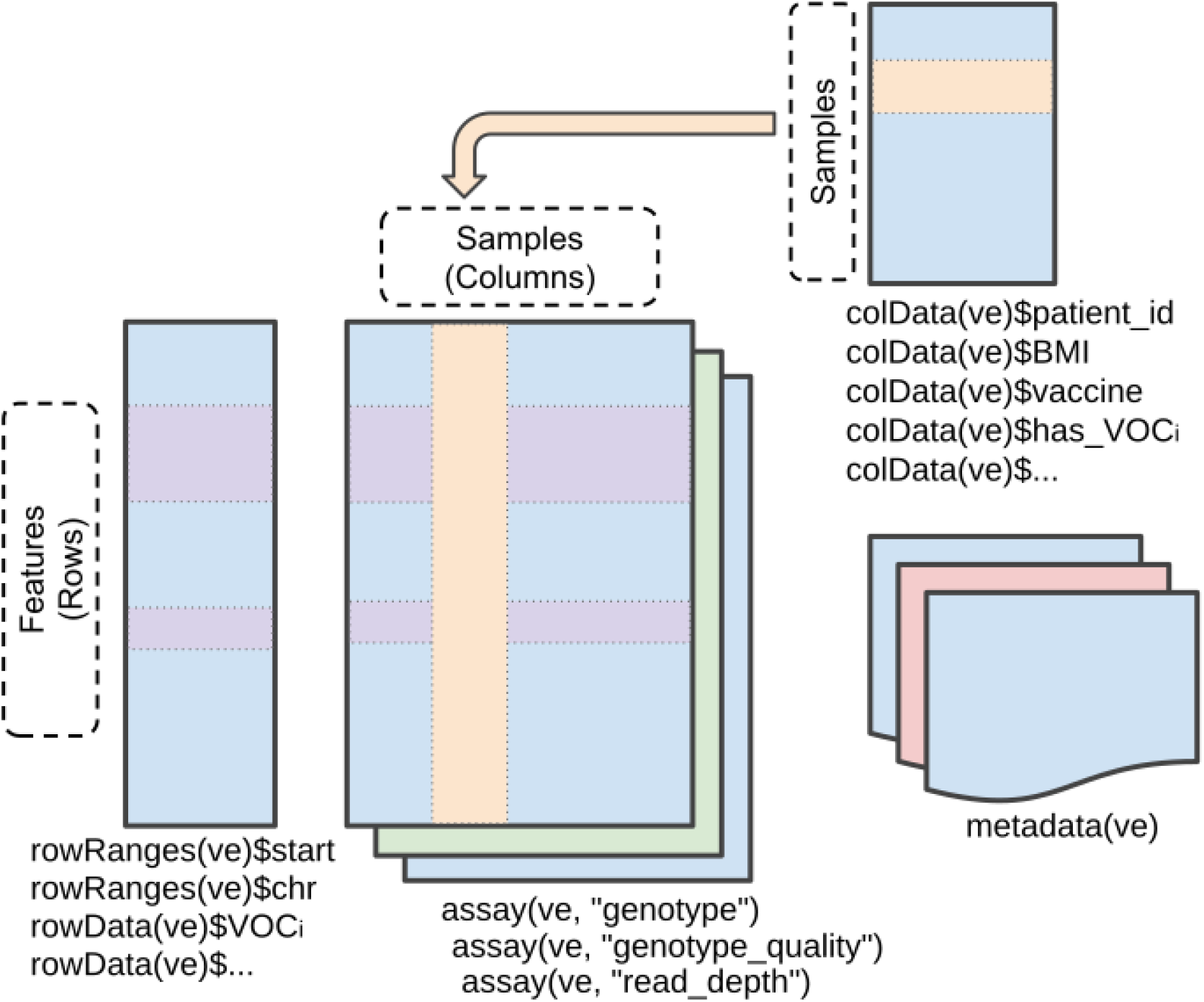
Illustration of a VariantExperiment object for handling SARS-COV2 genomic data and associated metadata. The latter should be interpreted in the broad sense of the term and includes sample characteristics, pre-analytical, analytical and data processing steps (stored in the colData component) as well as mutation and variant characterisation (stored in the rowData component). The generation and manipulation of such an object is demonstrated in our accompanying vignette [8]. The figure is adapted from the SummarizedExperiment package with permission of the author, Jim Hester.

### Genomic data

The VariantExperiment object model enables to store .vcf and .gds data into a structured data model, namely RangedSummarizedExperiment object [9]. These objects contain one or more assays, each represented by a matrix-like object. In the context of the SARS-CoV-2 genomic data, the first assay should therefore contain the called genotype (Figure 2, left). Additional assay can be used to store quality metrics associated with the genotype calling (e.g. genotype quality calculated as a Phred-scaled probability of the called genotype, read depth on Figure 2, right). It is worth noting that large assays can also be stored on disk (rather than in memory), thus enabling the handling and processing of large datasets (i.e., hundreds of variants and thousands of patients) that would not fit in memory.

**Figure 2:**
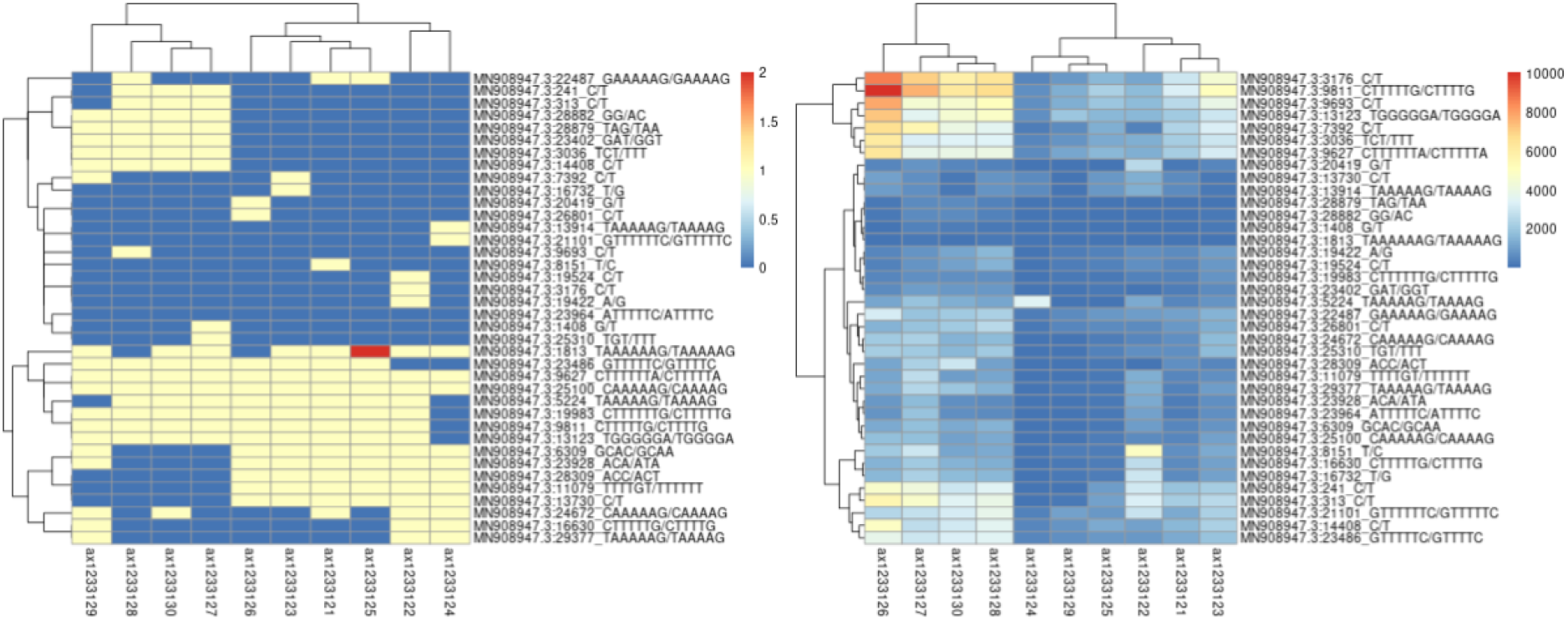
Visualisation of the two assay slots stored in our example data. On the left, the genotypes, encoded as 0s for the reference allele, and 1, 2,.. for alternative references. On the right, we see the sequencing read depth and how some samples have systematic low read depth (blue).

### Metadata

The metadata terms includes a wide range of data annotations, including sample characteristics, as well as variant and mutation characterisation. Linking genomic data and their associated metadata is crucial in omics sciences. Accordingly, international collaboration should be put in place in order to standardize the metadata ontology.

When using the VariantExperiment infrastructure, sample and feature metadata are saved as tabular data (DataFrame or DelayedDataFrame, when the data is large and requires on-disk access). The sample metadata, referred to specifically as colData (column/sample metadata) offers a structured way to describe sample characteristics (such as patient identifiers, age, any clinical variables such as BMI, or any comorbidities, whether patients were vaccinated, with which vaccine and when), pre-analytical and analytical parameters (such as library preparation or the sequencing platform that was used to generate the data), as well as indications about the specific bioinformatics pipelines (algorithms and software versions) that were used for batches of samples of various origins. The individual rows (features) are also annotated using exact genomics coordinates [10], protein-level alteration, as well as variants specification (e.g.VOC; see below) to cite a few. These annotations are referred to as *rowData* or *rowRanges* (when referring to genomic coordinates). As for the genomic data, that rapid accumulation of annotations will become a computational bottleneck, the on-disk representation of the metadata can be used to reduce its memory footprint, enabling the processing of many large files.

Another key aspect of this data model is the coordination of the assays and their row and columns metadata during all data manipulation, subsetting and processing. Accordingly, any subsetting along the assay rows (features) or columns (samples) are propagated along the colData and rowData/rowRanges metadata in a coordinated fashion. This property therefore ensures to keep a consistent synchronization between genomic data and all associated metadata.

### Mutation and variant characterization

As illustrated in Figure 1, the description of each called mutation is saved and annotated in the rowData and rowRanges metadata slots. In the context of the analysis of SARS-COV2 genomic data, this component should include several columns including the mutation position along the virus’s genomic coordinates as well as nucleic acid and amino acid changes. Moreover, an indicator may be inserted to reflect if a mutation is part of VOC or VOI signature, and should regularly be updated according to the evolution of scientific knowledge. One such indicator column should be added for each VOC and VOI (Figure 3).

**Figure 3:**
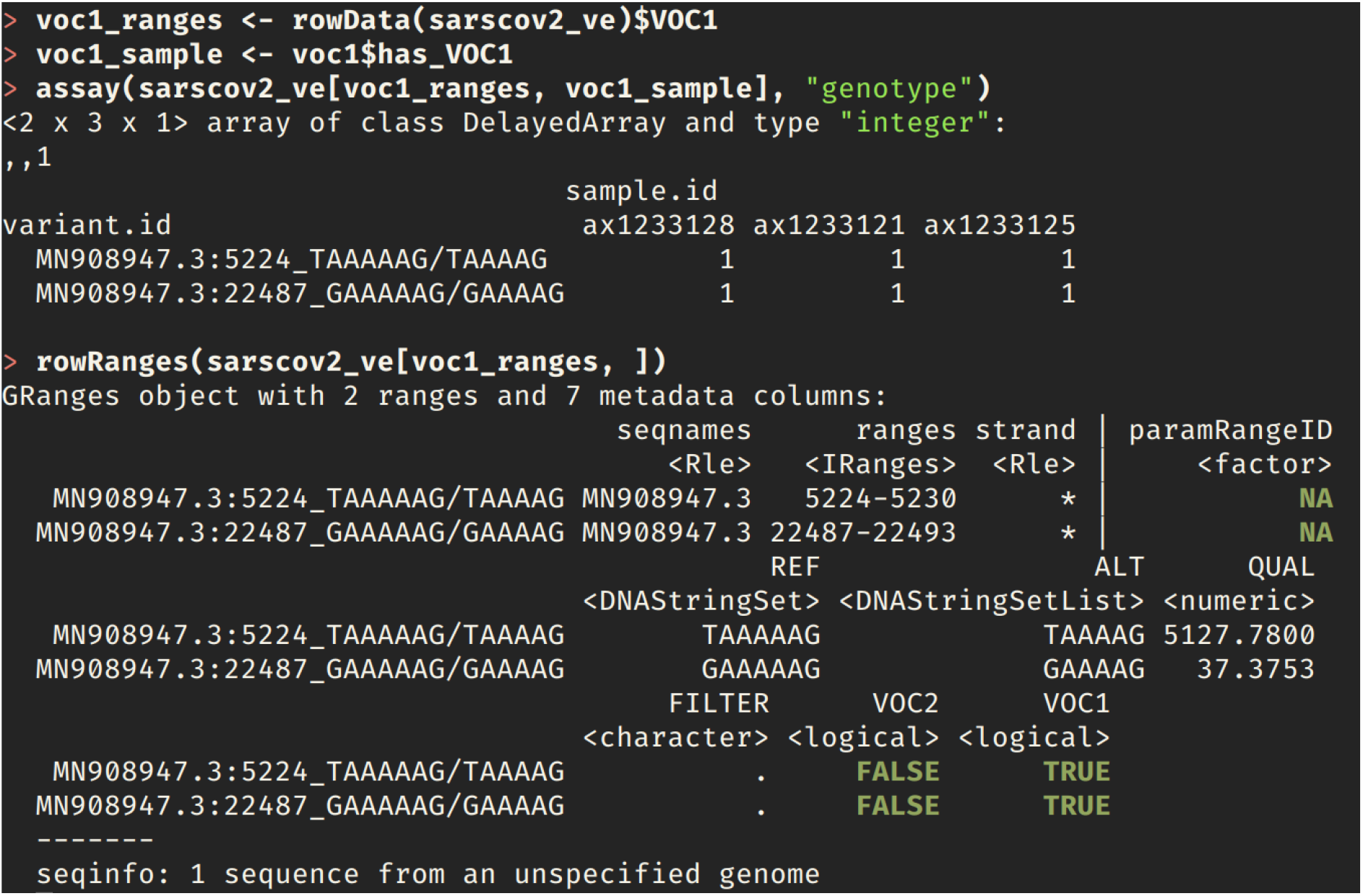
Illustration of the definition of a variant of interest (VOC1) and how the VOC1 indicator is used to extract the alleles that define it (voc1_ranges variable). The voc1_sample variable selects the samples that have been annotated as being infected by that variant. We then show how the 3 samples have the alternative allele (a deletion) in the 2 sites at positions 5224-5230 and 22487-22493 respectively.

On Friday 26th March WHO organized a meeting focusing on sero epidemiology for SARS-CoV-2 VOC. During this meeting WHO already reported 3 and 6 confirmed VOC and VOI, respectively. However, 16 other signals of potential VOIs/VOCs are in assessment. Accordingly, the number of columns in the rowData metadata slots is expected to grow in the following months/years. Conversely, samples can be annotated as having been infected by a particular VOC or VOI in the colData component (Figure 3). Examples of such variant and infection characterisation are documented in our companion vignette.

## Conclusions

In this paper, we illustrate how the VariantExperiment Bioconductor package can be used to store, share and analyze SARS-COV2 genomic data and their associated metadata. For many years, the Bioconductor infrastructure has proved to match the requirements for handling omics data (e.g. bulk or single-cell transcriptomics, genomics, proteomics,….). To this end, Bioconductor employs a flexible object-oriented paradigm that enables to encapsulate multiple object components into a single instance and preserve the relations between primary (i.e. genomic, transcriptomic,..) data and metadata. As proposed here, the integration of SARS-CoV-2 genomic data and associated metadata in the VariantExperiment object model stenghtens the discovery and annotation of new VOCs and VOIs among the fast expanding list of newly emerging SARS-CoV-2 variants. Moreover, the VariantExperiment object may include other complementary data such as quality metrics and technical parameters, among which pre-analytical (i.e library preparation kit), analytical (sequencing technology) and bioinformatics (software version and parameters) data. Integrating the latter into the same object would substantially broaden the field of genomic data exploitation, e.g. enable stakeholders to answer queries that address the impact of technical parameters on data quality.

## Footnotes

1 https://uclouvain-cbio.github.io/VariantExperiment-COVID19-UseCase/

2 https://github.com/UCLouvain-CBIO/VariantExperiment-COVID19-UseCase

